# The domed architecture of *Giardia*’s ventral disc is necessary for attachment and host pathogenesis

**DOI:** 10.1101/2023.07.02.547441

**Authors:** KD Hagen, C Nosala, NA Hilton, A Müller, D Holthaus, M Laue, C Klotz, A Aebisher, SC Dawson

## Abstract

After ingestion of dormant cysts, the widespread protozoan parasite *Giardia lamblia* colonizes the host gastrointestinal tract via direct and reversible attachment using a novel microtubule organelle, the ventral disc. Extracellular attachment to the host allows the parasite to resist peristaltic flow, facilitates colonization and is proposed to cause damage to the microvilli of host enterocytes as well as disrupt host barrier integrity. The 9 µm in diameter ventral disc is defined by a highly complex architecture of unique protein complexes scaffolded onto a spiral microtubule (MT) array of one hundred parallel, uniformly spaced MT polymers that bend approximately one and a quarter turns to form a domed structure. To investigate the role of disc-mediated attachment in causing epithelial cell damage, we used a new approach to rapidly create a stable quadruple knockout of *Giardia* of an essential ventral disc protein, MBP, using a new method of CRISPR-mediated gene disruption with multiple positive selectable markers. MBP quadruple KO mutant discs lack the characteristic domed architecture and possess a flattened “crescent” or horseshoe-shaped conformation that lacks the overlapping region, with severe defects in the microribbon-crossbridge (MR-CB) complex structure. MBP KO mutants are also unable to resist fluid flow required for attachment to inert surfaces. Importantly, MBP KO mutants have 100% penetrance off positive selection, which is essential for quantification of *in vivo* impacts of disc and attachment mutants with host cells. Using a new gastrointestinal organoid model of pathogenesis, we found that MBP KO infections had a significantly reduced ability to cause the barrier breakdown characteristic of wild-type infections. Overall, this work provides direct evidence of the role of MBP in creating the domed disc, as well as the first direct evidence that parasite attachment is necessary for host pathology, specifically epithelial barrier breakdown.

## Introduction

*Giardia lamblia* is an anaerobic protistan parasite that causes the diarrheal disease giardiasis in both humans and animals [1]. Transmission is thought to be zoonotic, with the same strains able to infect different animal hosts. The disease disproportionately impacts regions of the world with poor sanitation and water quality and can be particularly severe in children and the immunocompromised [1]. Infections begin when mammalian hosts ingest environmentally resistant cysts from contaminated water or food sources. In the stomach, the cysts excyst into flagellated trophozoites that attach extracellularly to the gastrointestinal epithelium of the proximal small intestine using the ventral disc, a unique cup-like microtubule organelle (reviewed recently in [2]). Attachment presumably allows the parasite to resist peristaltic flow and facilitates colonization and proliferation through the close association of the parasite with the enterocytes of the proximal small intestine [1]. The bipartite *Giardia* life cycle is completed when trophozoites differentiate into cysts, which are shed in the feces and infect new animal hosts. Acute and chronic giardiasis include common symptoms of nausea, abdominal cramps, gas, and weight loss. Severe cases also include malabsorptive diarrhea with fatty, bulky stools [1]. Both malnutrition and delayed development have been associated with early and recurrent childhood giardiasis [1].

With over 300 million cases annually, giardiasis remains a “neglected disease” due to its worldwide prevalence and to a general lack of molecular understanding of both parasite biology and interactions with the host [3]. The parasite neither produces a known toxin nor induces a robust inflammatory response, yet at the cellular level, parasite colonization of the gut epithelium is associated with malabsorption, shortened microvilli, disruption of the intestinal barrier function, induction of enterocyte apoptosis, inhibition of brush-border disaccharidases and lactases, mucus degradation, loss of mucosal surface due to increased crypt to villus ratios, and manipulation of host immune responses via arginine/nitric oxide limitation [4]. *Giardia* colonization and proliferation in the small intestine also disrupts the ecological homeostasis of gastrointestinal commensal microbes [5]. Despite *Giardia’s* significance in global health, the mechanisms of pathogenesis are poorly understood and have been vaguely described as “multifactorial” [1] yet may include parasite metabolism, parasite “fitness”[6], secretion of cysteine proteases [7], or host microbiome dysbiosis [5].

An obvious, yet often overlooked contributor to the host cellular pathology observed in giardiasis is *Giardia’s* ventral disc [2]. Disc-mediated attachment of *Giardia* to the microvilli could directly induce cellular pathology in the host, yet this is perhaps the least explored potential pathogenic mechanism. Early observations of *Giardia* trophozoites attached to the epithelium showed prominent ventral disc-shaped depressions that remained in the microvilli after parasites had detached [5–7]. Disc- and lateral crest-mediated attachment by a single trophozoite would affect roughly 2000 microvilli, and attachment has been proposed to cause both direct and/or indirect host cellular damage that includes shortened microvilli and malabsorption.

The ventral disc is a distinctive and complex microtubule organelle that is unique to *Giardia* species [11]. The precise mechanism of disc-mediated attachment remains controversial [2], yet parasites readily and reversibly attach not only to the host epithelium, but also to inert laboratory surfaces such as coverslips or culture tubes. Approximately 9 µm in diameter, the disc is defined by a highly complex architecture scaffolded onto a spiral microtubule (MT) array of one hundred parallel, uniformly spaced MT polymers that curve approximately one and a quarter turns to form a domed structure [12]. The disc MTs are highly decorated with conserved and novel microtubule associated proteins (MAPs), as well as with novel disc-specific protein complexes such the “microribbon-crossbridge” complex [13] [14] and the “sidearm” and “paddle” complexes [15]. An additional fibrillar structure, the “lateral crest”, surrounds the periphery of the ventral disc [10,16].

The lack of direct evidence linking disc-mediated attachment to cellular damage is due in part to the limited access to host sites of infection and the lack of genetic tools to evaluate the functioning and specific attachment defects of mutant discs [17]. Our lab has facilitated a molecular-based understanding of the ventral disc through the development and use of new molecular genetic tools to examine disc structure and function [17]. We have also identified close to 90 disc-associated proteins (DAPs) that localize to particular structural regions of disc, including specific DAP subsets that localize exclusively to the overlap zone, lateral crest, disc margin, or ventral groove (Figure 1 and [2]). Using morpholino and CRISPRi-based knockdowns, we have also previously shown that several novel DAPs are associated with MT and overall disc hyperstability [18], as well as disc architecture [19,20].

**Figure 1.**
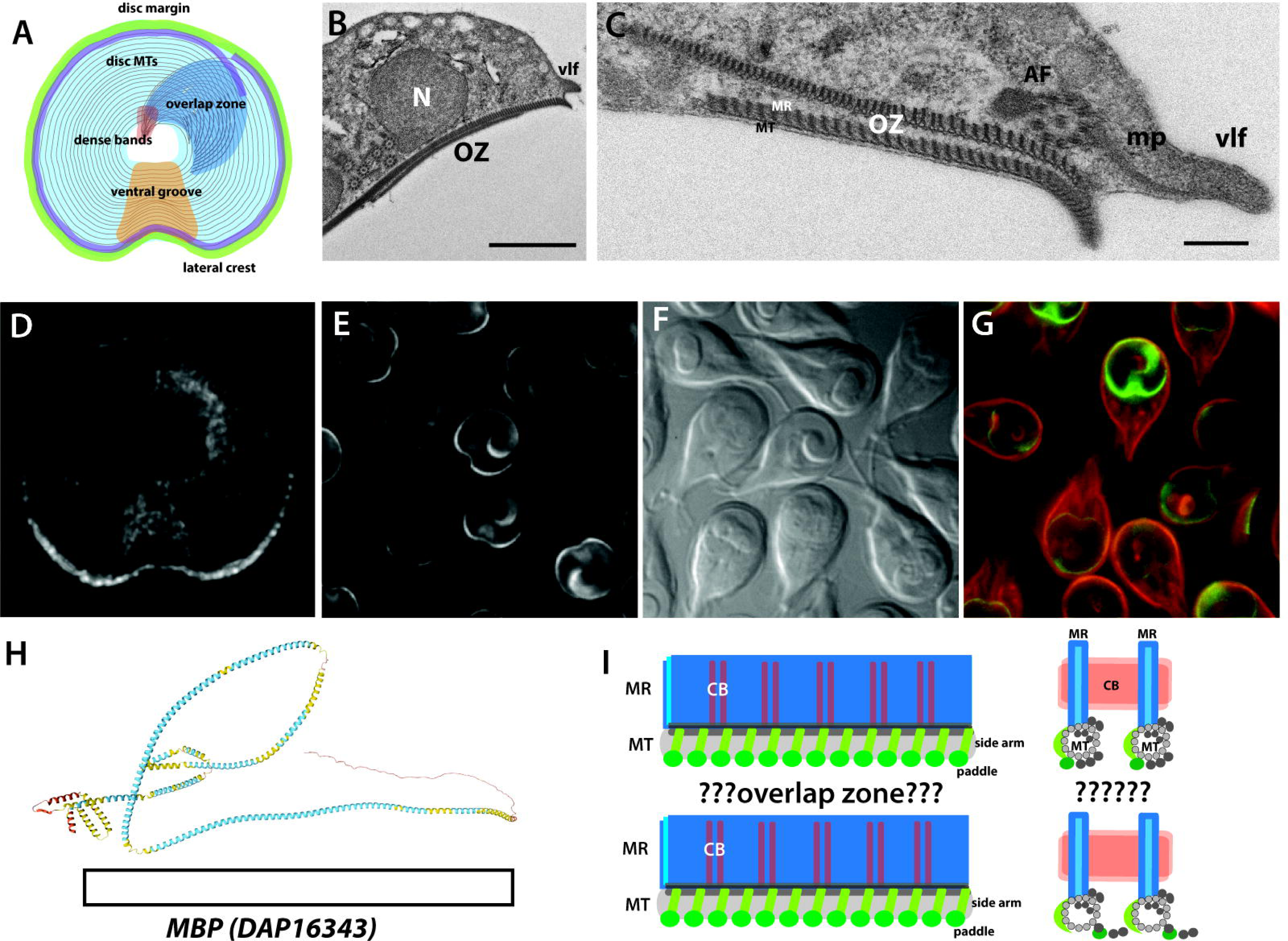
Novel DAP16343 (median body protein, MBP) localizes to the disc body and overlap zone. Distinct structural regions of the disc [2]and lateral crest (LC) are noted in the schematic, which includes the disc body (body), the ventral groove (vg), the disc margin (dm), the overlap zone (oz), and the dense bands (db). The overlap zone (oz) region is an area of the disc close to the disc margins highlighted in the TEM cross-section in B, C. The lateral crest (LC, green) is an additional non-MT, yet repetitive structure surrounding the outer edge of ventral disc [2]. C-terminally GFP-tagged MBP primarily localizes to the overlap zone, disc margin, and disc body in SIM (D) and conventional epifluorescence images (E, F). In G, the MBP-GFP (green) fluorescence is shown in relation to the cell membrane (CellMask, red). In H, the predicted AlphaFold protein structure highlights the lack of resolution of the DAP16343 (MBP) which lacks homology to proteins outside of the *Giardia* genome. While the ventral disc microribbons (MR) are associated along the length of the ventral disc MTs (MT) in both upper and lower portions of the overlap zone (I), it remains unclear as to how upper and lower portions interact or are connected.

Here we advance our understanding of the role of disc-mediated attachment in causing epithelial cell damage using a new approach to rapidly create a stable knockout of MBP, a known disc protein of unknown function, by the insertion of multiple positive selectable markers. MBP KO discs have a flattened “crescent” or horseshoe-shaped conformation that lacks an overlap zone and is structurally similar to knockdown phenotypes we have previously observed using morpholinos or CRISPRi-based methods [19,20]. Due to severe defects in microribbon height and spacing, we predict that the MBP protein is a component of the MR-CB complex. Using MBP KO mutants, we confirm the role of disc doming in proper trophozoite attachment under fluid flow and of the lateral crest in establishing and maintaining membrane contacts that form a “seal” with the attachment surface. MBP KO mutants have 100% penetrance off positive selection, which is essential for quantification of *in vivo* impacts of disc and attachment mutants with host cells. Using new gastrointestinal organoids, we determine that barrier breakdown is dramatically reduced in infections with MBP KO mutants. Thus, for the first time, we provide direct evidence to show that parasite attachment is necessary for host epithelial barrier breakdown.

## Materials and Methods

### Giardia cultivation, electroporation, and isolate cloning

*Giardia lamblia* trophozoites (strain WBC6, ATCC 50803) were grown at 37°C without shaking in 16-ml screw-cap plastic tubes (BD Falcon) containing 12 ml modified TYI-S-33 medium supplemented with bovine bile, 5% adult and 5% fetal bovine serum. Upon reaching confluency, or after 48 hrs (MBP KO), tubes were incubated on ice for 30 minutes and were subcultured by transferring detached trophozoites (0.5 to 1 ml) to warmed medium [21].

Electroporations of plasmids or linear homology-directed repair (HDR) templates into *Giardia* were performed as described previously [18], using 20 to 40 μg DNA and 1 x 10^7^ trophozoites per electroporation. Electroporated cells were transferred to standard culture tubes and incubated at 37°C. Medium was replaced every 48 hours. Lower antibiotic concentrations were used for initial selection (see below) and were increased when trophozoites reached >50% confluency.

Dilution to obtain clonal cell lines was performed in 24 well tissue culture plates (Corning). Three rounds of dilution, first to 10 cells/well, and subsequently to 0.8 cells per well, were performed. Plates were incubated at 37°C in Mitsubishi AnaeroPack 2.5L rectangular jars with a Mitsubishi AnaeroPack-Anaero Gas Generator sachet (ThermoScientific) for 7 to 10 days, at which point wells were examined for growth. Trophozoites were detached from wells by incubating on ice for 30 minutes. Aliquots were retained for PCR screening (see below), and the remainder was transferred to standard culture tubes for further propagation.

### Design and evaluation of Giardia-optimized hygromycin, blasticidin, and neomycin drug resistance markers

Modern genetics in tetraploid *Giardia* requires effective selectable markers. Two antibiotic resistance markers, pac and neo, conferring resistance to puromycin and G418, respectively, have been used in *Giardia* for more than 25 years [22,23]. A third marker, bsr (from *Bacillus*; blasticidin) has been used infrequently [24,25]. Anticipating a need for additional robust markers, we tested the sensitivity of wild type WBC6 to G418 (100 to 400 μg/ml), blasticidin (75 to 300 μg/ml), hygromycin B gold (300 to 1200 μg/ml), and phleomycin (10 to 500 μg/ml) (Supplemental Figure 2). Standard culture conditions were used (see above), and tubes were inoculated with 5 x 10 trophozoites. Cultures were examined microscopically every 24 hours for 72 hours and confluence was estimated. All antibiotics were purchased from Invivogen except G418 (Fisher Bioreagents) and all but phleomycin were lethal to *Giardia* at one or more of the concentrations tested.

New antibiotic cassettes were designed to maximize antibiotic resistance gene expression in *Giardia* (Supplemental Figure 1). Antibiotic resistance gene sequences from vectors pHJ41 (BSD from *Aspergillus*, blasticidin) [26], pIR3HYG (HYG, hygromycin) [27], and pKS_mNeonGreen-N11_NEO (NEO, neomycin) [28] were codon-optimized for *Giardia* using the codon usage calculator %MinMax [29] with a codon usage table generated from the HIVE-CUTs database [30]. 200 bp fragments containing promoter and termination sequences from highly expressed *Giardia* ribosomal proteins [31] were added to the 5’ and 3’ ends, respectively, of the resistance genes. Unique promoter and termination sequences were used for each cassette to minimize the possibility of unwanted recombination between markers. The *Giardia*-optimized cassettes BsdRgo, HygRgo, and NeoRgo were synthesized (Twist Bioscience) and cloned into pcGFP1Fpac [16], replacing GFP. Each plasmid was introduced into wild type *Giardia* by electroporation (see above) and selection with puromycin. The antibiotic sensitivity of strains bearing the AbxRgo marker plasmids was then tested (Supplemental Figure 1). Optimal drug concentrations for initial and final selection of plasmids with the new cassettes were: G418 (150 µg/ml and 600 µg/ml), blasticidin (75 µg/ml and 150 µg/ml), and hygromycin B gold (600 µg/ml and 1200 µg/ml. The antibiotic concentrations listed here were also used to select for stably integrated HDR-Abx cassettes when knocking out MBP alleles.

### Design of the Cas9/gRNA expression vector and HDR templates for CRISPR based quadruple knockout of MBP (DAP16343)

Our CRISPR strategy for MBP knockout employed a Cas9/gRNA expression vector, Cas9g1pac, that is nearly identical to our previously published *Giardia* CRISPRi vector dCas9g1pac [17], except that the mammalian codon optimized SpCas9 from JDS246, a gift from Keith Joung (Addgene plasmid # 43861; http://n2t.net/addgene:43861), lacks the D10A and H840A mutations that render dCas9 inactive. As in our CRISPRi vector, Cas9 expression is driven from the *Giardia* malate dehydrogenase (MDH, GL50803_3331) promoter, and the sequences C-terminal to Cas9, including the native *Giardia* GL50803_2340 NLS that directs Cas9 to the nuclei, are unchanged. As previously described, gRNA expression is driven by the *Giardia* U6 spliceosomal snRNA promoter, and an 18 bp region containing inverted BbsI sites for cloning annealed gRNA oligos lies between the U6 promoter and the gRNA scaffold. For MBP knockout, a gRNA (20 nt) targeting MBP (16343gRNA95) was designed as previously described [17] using the CRISPR ‘Design and Analyze Guides’ tool from Benchling (https://benchling.com/crispr) with an NGG PAM sequence and the *Giardia lamblia* ATCC 50803 genome (GenBank Assembly GCA_000002435.1). Forward and reverse gRNA oligos (5’-caaaAACATTTGCTGAGCAGGTCA-3’ and 5’-aaacTGACCTGCTCAGCAAATGTT-3’) including 4-base overhangs complementary to the vector sequence overhangs were annealed and ligated to BbsI-digested Cas9g1pac to create the CRISPR expression vector Cas9MBPgRNA95.

HDR templates for MBP knockout consisted of the 750 bp immediately up- and downsteam of the DSB site (left and right homology arms), plus one of the three *Giardia*-optimized antibiotic cassettes (see above and Supplemental Figure 2). Mutations made within the gRNA binding site ensured that repair templates were not cut by Cas9. Homology arm fragments were synthesized (Twist Biosciences) and cloned into the SacI and KpnI sites of pBluescriptII KS (+). Antibiotic cassettes were then inserted at a unique BamHI site created at the DSB site. Linear HDR templates for knockout were PCR amplified using Phusion DNA polymerase (New England Biolabs) with T7 Forward and M13 Reverse primers. PCR product was purified with Zymo Research Clean and Concentrator-25 columns.

### Creation of the MBP (DAP16343) quadruple knockout strain

For gene knockout, a strain expressing Cas9 and the specific gRNA is required. To generate the expression strain for MBP knockout, 20 µg of the CRISPR expression vector Cas9MBPgRNA95 was electroporated into wild type WBC6 [32], as described above. The expression strain was maintained with puromycin selection (50 µg/ml) throughout the knockout process, Linear Abx HDR templates were introduced sequentially by electroporation. BsdRgo was introduced first, followed by NeoRgo, and then HygRgo. After each round of electroporation, total DNA was extracted from cells using a Zymo Research Quick DNA Midiprep kit, and PCR was performed to screen for the presence of a larger fragment indicating the new marker using primers MBPLeftF (5’-GCGCTTAAGCTCAATGGTCG-3’) and MBPRightR (5’-TCTGCTCGCTCTAGAGCCTC-3’), which bind outside of the HDR template homology arms. After introduction of the third cassette, dilutions of the MBP triple KO were performed in 24 well tissue culture plates (see above) to screen for MBP quadruple KOs. After the first round of dilution, a cell population with fewer copies of wild type MBP was found (Supplemental Figure 3). After the second round, quadruple KO clones 9A, 9F and 9I, that lacked a wild type copy of MBP were discovered among the triples (Supplemental Figure 2). A third round was performed to ensure clonal cell lines. All MBP quadruple KO strains were thawed from frozen stocks and cultured for at least 48 hours prior to phenotypic analysis, unless otherwise specified.

### Immunostaining of MBPKO and other tagged DAP strains

Prior to immunostaining, *Giardia* trophozoites grown to confluency were either fixed and settled on coverslips or were attached live to coverslips and then fixed. For settled trophozoites, 4% paraformaldehyde, pH 7.4, was added to culture tubes (37°C, 15 minutes). Trophozoites were then pelleted at 900 x g for five minutes, washed three times with 2 ml PEM, pH 6.9 [33], incubated in 0.125M glycine (15 minutes, 25°C), and washed three more times with PEM. Fixed cell suspensions were then settled on coverslips coated with poly-L-lysine. Cells attached live were iced 30 minutes, harvested, washed three times in 1x HBS, pH 7.0 and resuspended in 250 µl HBS. Aliquots were added to warm coverslips (15 minutes, 37C) and were then fixed *in situ* with 4% paraformaldehyde, pH 7.4 for 15 minutes, washed three times with 2 ml PEM, and incubated in 0.125M glycine (15 minutes, 25°C). Both attached and settled trophozoites were washed again three times with PEM, prior to permeabilization with 0.1% Triton X-100 for 10 minutes. Three PEM washes were performed and cells were blocked in 2 ml PEMBALG for 30 minutes [20]. Coverslips were incubated overnight at 4°C with anti-delta-giardin (1:1000) or anti-beta-giardin (1:1000, both gifts of M. Jenkins, USDA) or with anti-TAT1 antibodies (1:250). Coverslips were washed 3 times in PEMBALG and incubated with goat anti-rabbit and/or goat anti-mouse Alex Fluor 488, 594, or 647 antibodies (1:1000; Life Technologies) for four hours at room temperature. Following incubation with the secondary antibodies, coverslips were washed three times each with PEMBALG and PEM and mounted in Prolong Diamond antifade reagent (Invitrogen). All imaging experiments were performed with three biologically independent samples.

### Super-resolution SIM 3D imaging of fixed cells

For super-resolution imaging of fluorescently-tagged DAPs or immunostained MBP KO strains, three dimensional stacks were collected at 0.1 µm intervals using a Nikon N Structured Illumination Super-resolution Microscope (SIM) with a 100x/NA 1.49 objective, 100 EX V-R diffraction grating, and an Andor iXon3 DU-897E EMCCD. Images were collected and reconstructed using NIS-Elements software (Nikon) in the “3D-SIM” mode (with no out of focus light removal; reconstruction used three diffraction grating angles each with three translations). Images were reconstructed in the “Reconstruct Slice” mode and were only used if the reconstruction score was 8. Images are displayed as a maximum intensity projections or 3D stacks.

### Quantification of ventral disc doming

To quantify disc curvature, trophozoites were attached to imaging dishes (MaTtek) and incubated at 37°C for 30 minutes in 1x HBS, washed three times with 1X HBS to remove unattached cells, and resuspended in 100 µl of warm 1x HBS. For live imaging, an equal volume of warmed (37°C) 3% low melt agarose in 1X HBS was added to immobilize trophozoites (final concentration 1.5%). Cells were maintained at 37°C during disc imaging. Images were acquired with a Leica DMI 6000 wide-field inverted fluorescence microscope using Metamorph acquisition software (Molecular Devices). Optical slices were collected at 0.2 µm intervals up to 8 µm stacks to capture the entire disc. Disc doming images were generated by reslicing the stack laterally across the posterior portion of the bare area; angles were measured using ImageJ/Fiji [31]. For each experimental condition, at least 30 cells were measured on three separate days totaling 90 cells imaged and quantified.

To evaluate the effects of drugs targeting microtubule dynamics on disc doming, a final concentration of 20 µM Taxol or 10 µM nocodazole was added to culture medium for one hour or five hours, respectively, prior to immobilization and imaging of cells. For disc doming measurements at 4°C, parasites were first immobilized in agarose, then the imaging plates were incubated on ice for 30 minutes prior to imaging. Live cells were fixed by adding ice cold 1x HBS containing a final concentration of 4% paraformaldehyde to cells incubated on ice for 15 minutes. Fixed cells were then pelleted, washed three times in chilled 1x HBS, and imaged in a pre-chilled coverglass bottom 96-well plate as previously described.

To image parasites attached to the MCF 10a human epithelial cell line, MCF10a cells were grown in a 96-well imaging plate in a 5% CO2-humidified incubator at 37°C in Complete MCF-10A Growth Medium composed of DMEM/F12 (Mediatech, Inc., Herndon, VA) supplemented with 5% donor horse serum, 20 ng/ml epidermal growth factor (EGF), 10 μg/ml insulin, 0.5 μg/ml hydrocortisone (Sigma, St. Louis, MO), 100 ng/ml cholera toxin (Cambrex, Westborough, MA), and 100 units/ml penicillin and 100 μg/ml streptomycin (Invitrogen, Carlsbad, CA). Prior to imaging with the *Giardia* 86676GFP (delta-giardin) strain, MCF10a cells were washed three times with warmed 1x PBS and incubated in 180 µl 1x PBS. Twenty microliters of chilled trophozoites in 1x PBS (100,000 cells) were added to each well. This co-culture was incubated at 37°C for 15 minutes to allow parasite attachment. Wells were washed three times in warm 1x PBS to remove unattached parasites before imaging.

For imaging isolated discs, the detergent extraction of the entire microtubule cytoskeleton was performed as previously described (Hagen et al., 2011) with some modifications. Briefly, *Giardia* medium was decanted from each confluent culture tube and trophozoites were washed three times with 5 ml of warmed 1x HBS. To demembranate cytoskeletons, cells were incubated on ice for 15 minutes in the last 1x HBS wash, pelleted by centrifugation (1000 × g, 5 minutes), and resuspended in 1 ml of 1xPHEM (60 mM PIPES, 25 mM HEPES, 10 mM EGTA, 1 mM MgCl2, pH 7.4) containing 1% Triton X-100 and 1M KCl. This solution was transferred to a 1.5 ml Eppendorf tube and vortexed continuously for 30 minutes at medium speed. Ventral disc cytoskeletons were pelleted at 1000 x g for five minutes and washed twice in 1X PHEM lacking both Triton X-100 and KCl. Sufficient extraction of cytoskeletons was confirmed by wet mount using DIC microscopy. Extracted cytoskeletons in PHEM were added to the 96-well glass bottom imaging plate and maintained at 37°C or on ice for 30 minutes prior to imaging as described.

Disc and bare area dimensions including perimeter and area, angle of repose (doming), and surface contact dimensions were quantified from > 300 immunostained images per condition using Fiji [34]

### Imaging of cell surface contacts of live attached trophozoites

To image cell surface contacts, trophozoites were harvested, washed once with 5 ml 1x HBS, pH 7.0 and resuspended in 5 ml 1x HBS. CellMask Orange (Invitrogen) was added to a final dilution of 1:5000. Cells were incubated on ice 5 minutes and then washed three times in 5 ml 1x HBS. Cells were added to 96-well #1.5 black glass bottom imaging plates (CellVis) and allowed to attach at 37C for 30 minutes. Single focal plane images at the attachment surface were acquired with a Leica DMI 6000 wide-field inverted fluorescence microscope using the 100x objective and μManager image acquisition software. Disc surface contacts were quantified and scored from >200 CellMask-stained images.

### Quantification of shear forces of using a flow cell assay of attachment

MBP knockouts, Cas9 control, and wild-type WBC6 trophozoites were iced 15 minutes, washed with 1x HBS and stained with CellMask Orange for 10 minutes. Cells were washed again with 1x HBS before addition to Ibidi mSlide VI 0.4 flow chambers to quantify their ability to resist shear forces from flow. Single focal plane images of trophozoites (2 million per ml) were acquired using µManager image acquisition software [35] with a 40X objective and a Leica DMI 6000 wide-field inverted fluorescence microscope. Epifluorescent images were collected of attached cells in chambers before and after a 20 second challenge with 3 ml/min flow for shear stress. Timelapse DIC images were collected at a rate of 1 per sec for 20 sec before challenge, during challenge, and 20 sec after challenge. Cells were allowed to attach for five minutes prior to initiation of flow. Attached cells were counted by overlaying the pre-flow epifluoresent image over the post-flow image (false colored green). Red cells (present pre flow only) were counted as unable to resist shear forces whereas yellow cells (present pre and post flow) were able to resist shear flow forces. The overall percent attached was calculated by quantifying the ratio of shear force resisting cells (yellow) to the total number of pre-flow cells (red and yellow).

### Human organoids and ODM-Giardia infections

Human organoids were generated and maintained from duodenal biopsy specimens obtained from healthy volunteers undergoing routine examinations at Charité-Universitätmedizin Berlin. The study was conducted with ethics approval (#EA4-015-13) from local authorities. The method for generating human organoids was previously described [36]. To culture organoid-derived monolayers (ODMs), Matrigel-coated transwell cell culture inserts (0.6 cm2, 0.4-mm pores; Corning, Tewksbury, NY) were used following established protocols [36].

For infection experiments, both wild-type and MBP quadruple KO trophozoites were passaged the day before to ensure logarithmic growth phase. Prior to infection, the parasites were detached from culture tubes by incubating them at 4°C for 20 minutes. The detached parasites were then pelleted and quantified using a Neubauer counting chamber. After quantification, the parasites were harvested by centrifugation at 1000 x g for five minutes at 4°C, followed by resuspension at the desired final concentration. To prepare for infection, the apical compartment medium of 8- to 10-day-old ODMs was replaced with complete TYI-S-33 the evening before the infection to allow the cells to adapt to the TYI-S-33 medium. Prior to infection, the medium was refreshed, with TYI-S-33 in the apical compartment and organoid differentiation medium in the basal compartment. *Giardia* trophozoites were added to the apical compartment and incubated for a duration of up to 72 hours.

### Transepithelial electric resistance (TEER) measurements of epithelial barrier integrity

To quantify barrier integrity in disc mutants, transepithelial electric resistance (TEER) measurements were performed using a Millicell ERS-2 Voltohmmeter (Merck-Millipore) equipped with an Ag/AgCl electrode (STX01; Merck-Millipore) on a 37°C heating block. The electric resistance of the blank (cell-free transwell insert) was subtracted from the raw resistance values and standardized for 1 cm2 surface area as previously described [36].

## Results

DAP16343 (median body protein, MBP) and the many other DAPs or MT-associated proteins that comprise the ventral disc have a structural role in creating or maintaining the disc domed architecture or have more specific functional roles in trophozoite attachment to inert surfaces or the gastrointestinal epithelium [11,18,33]. A key architectural feature of the spiral MT array of the ventral disc is a region of overlap, termed the “overlap zone” (OZ), where the upper and lower layers of the disc may interact (Figure 1A,B,C and I). MBP is an abundant disc protein [16] that localizes to both the disc body and the overlap zone (Figure 1D,E, and G). Like many DAPs, it lacks homology to known proteins in other organisms, and structural predictions (Figure 1H) provide little information to illuminate its role in the cell. Despite prior work using MBP knockdowns (morpholino or CRISPRi-based [17,33] it remains unclear how the domed disc architecture is achieved more generally, and how specific disc-associated proteins like MBP create, maintain, or modulate this domed architecture required for attachment.

### The ventral disc is domed at physiological temperature regardless of attachment surface or treatment

To facilitate attachment, the disc is proposed to maintain a rigid domed architecture resembling an inverted cup. To compare any differences in disc doming under various experimental conditions and temperatures, we used 3D imaging to quantify disc doming and measured the “angle of repose” (Figure 2) relative to the attachment surface in *Giardia* strains with fluorescent markers of either the ventral disc microribbons (dGiardin-GFP) or the ventral disc microtubule array (mNeonGreen-bTubulin)[18]. At physiological temperature (37°C), the average angle of repose was 16-17 degrees relative to an inert surface (coverslip) using either of the two disc markers (dGiardin-GFP or GFP-bTubulin) (Figure 2A,G). Because the rigidity of an inert surface may impact the degree of disc doming in attaching trophozoites, we also cultured parasites on a monolayer of MCF10A human epithelial cells (ATCC CRL-10317). Compared to the discs of trophozoites attached to inert glass surfaces, the average angle of repose was significantly increased (22 degrees) when trophozoites were attached to the biological and deformable MCF 10A epithelial cell membrane (Figure 2C,D,G). *Giardia*’s microtubule cytoskeleton, including the ventral disc and all eight flagella, can be readily extracted using detergent buffers, and has been imaged extensively using cryoEM [16, 17]. Detergent extracted, isolated ventral discs had significantly increased curvature at 37°C – with the average angle of repose at 25 degrees (Figure 2A).

**Figure 2:**
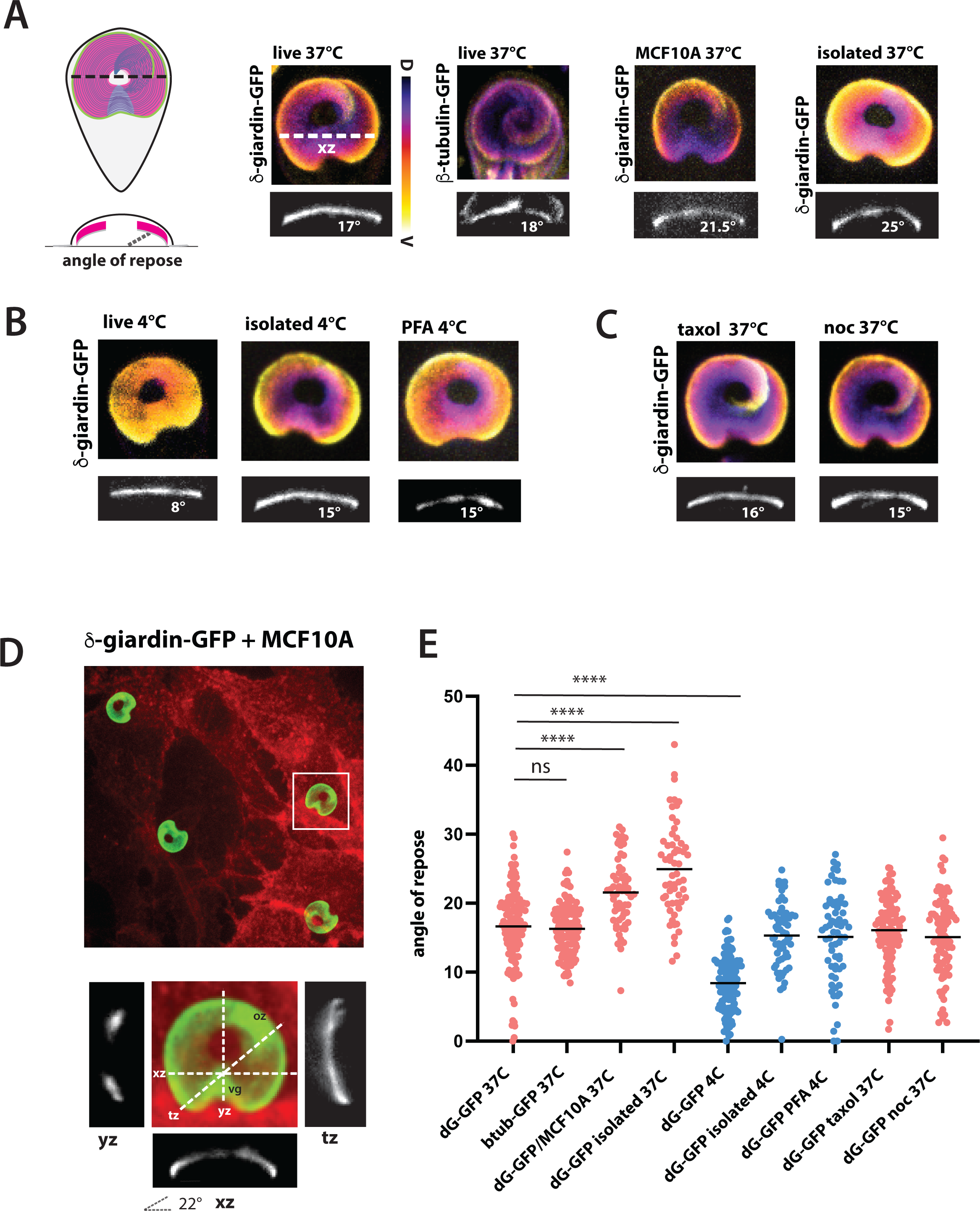
Marked disc doming is associated with *in vitro* and *in vivo* attachment. Using the DAP86676GFP C-terminal GFP tagged strain (dG-GFP) that marks the disc body [2] or a beta-tubulin GFP strain (btub-GFP), the overall angle disc doming (xz plane) was quantified with 3D imaging at variations in surfaces at 37°C (A), incubation at 4°C (B), or following treatment with drugs that disrupt microtubule dynamics (noc= nocodazole; taxol = paclitaxel) (C). Trophozoite attachment to the human epithelial cell line MCF10A (A, D), cytoskeletal isolation by detergent extraction (“isolated”A, B), or paraformaldehyde (B). Images are presented as 2D projections of 3D stacks and colorized using the temporal color code plugin in [34] from dorsal (purple) to ventral (orange) to highlight doming. Disc doming in three dimensions (xz, xy, yz) in d-G-GFP tagged parasites (green) attached to the human MCF10a cell line (red) is also presented in panel D. The disc doming angle relative to the surface was also quantified for over 50 replicate samples for each treatment (E). Triple asterisks indicate statistically significant differences (P< 0.001) in disc doming as compared to the untreated, tagged strains using the Mann-Whitney unpaired t-test (Methods).

*Giardia* trophozoites cultured at 37°C are commonly detached from culture tubes by incubation at 4°C for at least 15 minutes [32]. To evaluate disc curvature at 4°C in detached trophozoites, imaging dishes were placed on ice and immediately imaged. In cold detached trophozoites, the average disc angle of repose was significantly lower at 8 degrees as compared to 17 degrees at 37°C (Figure 2A, B,G). Chilled isolated discs had a higher angle of repose at 15 degrees, yet it was significantly lower than isolated discs at 37°C (Figure 2A). The addition of paraformaldehyde to ice cold trophozoites resulted in an angle of repose of 15 degrees that resembled discs incubated at 37°C without fixation (Figure 2B,G).

Lastly, while microtubule dynamic instability does not occur in interphase disc MTs [9, 33], presumably due to the MT stabilization by DAPs[18], other variations in MT polymerization dynamics during attachment could potentially impact the disc domed architecture. Treatment of attached parasites with either Taxol or nocodazole, drugs that limit microtubule dynamic instability, had no significant effect on disc angle of repose (16 and 15 degrees, respectively) as compared to the GFP tagged d-Giardin or b-Tubulin at 37°C (Figure 2C).

### The quadruple allelic knockout of DAP16343 (MBP) has severe and completely penetrant defects in disc architecture, including aberrant MR-CB complexes

We have previously shown that either morpholino or CRISPRi knockdown of MBP (DAP16343) in *Giardia* trophozoites results in a flattened disc with decreased angles of repose, and that these cells are also significantly decreased in their resistance to shear and normal forces of attachment [26, 30]. Here we used newly developed genetic tools to insert new antibiotic resistance markers that stably disrupt all alleles of the disc-associated protein DAP16343 (MBP), creating stable knockout (MBPquadKO) strains (Supplemental Figure 3). As seen with SIM, the MBPquadKO has a severely flattened disc with a low angle of repose (Figure 3A) reminiscent of that seen in the discs of chilled, detached trophozoites (Figure 2B).

**Figure 3:**
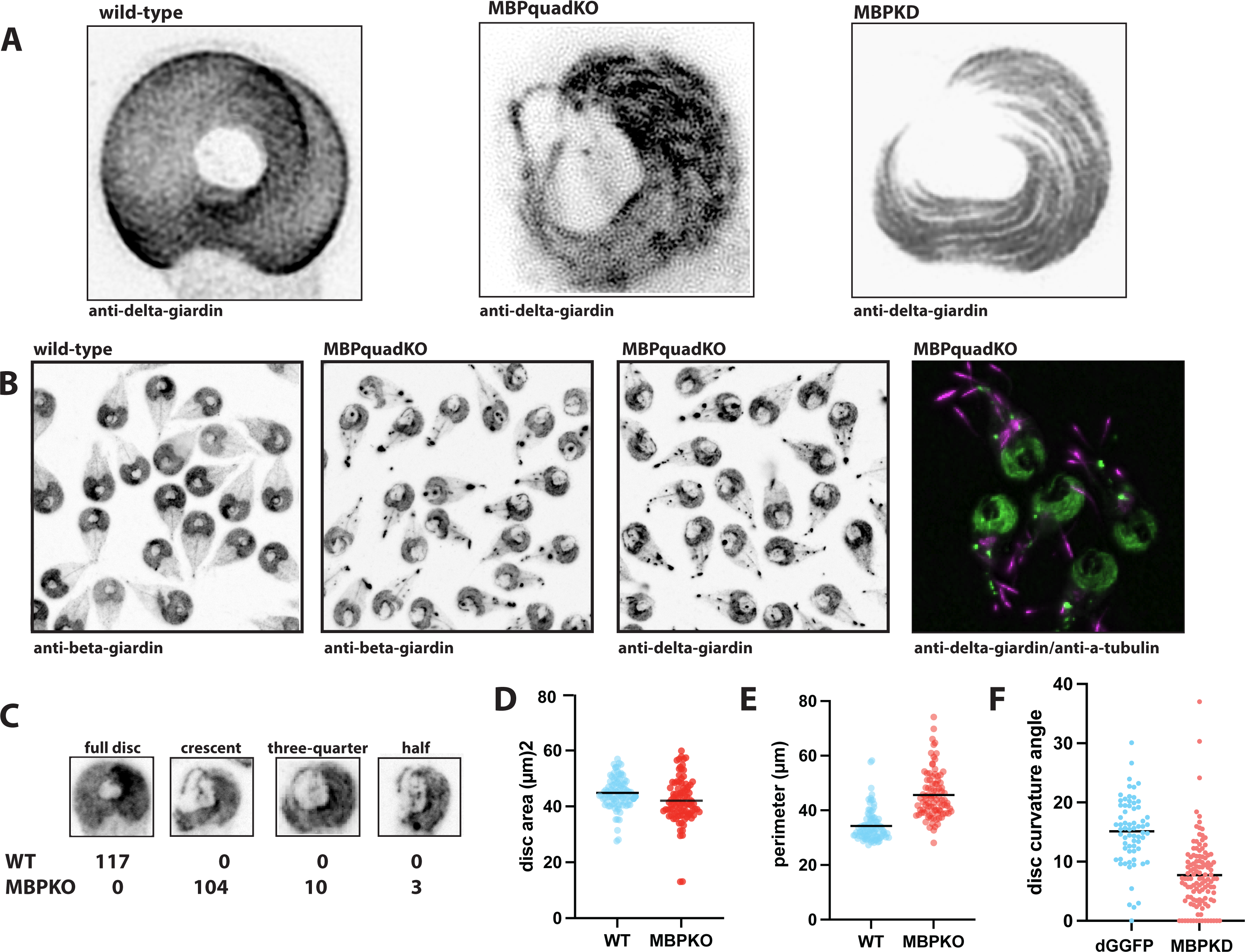
The DAP16343 (MBP) quadruple knockout mutant (MBPquadKO) has severely deformed and flattened discs with complete phenotypic penetrance. Structured illumination microscopy (SIM) imaging of the quadruple allele DAP16343 KO (MBPquadKO) or morpholino-based KD (A) in 3D demonstrates severely aberrant disc structure (Figure 2). As compared to wild-type, the MBPquadKO mutant (B) has severe defects with 100% penetrant aberrant discs (C) as seen by counterstaining with antibodies to the microribbon proteins beta- and delta-giardin. The majority (close to 90%) of MBPquadKO mutant discs have a “crescent” structure, yet up to 10% have either “horseshoe” or “half-moon” structures that are defined by the lack of an overlap zone (C). Mutant discs have similar overall surface areas (D), yet mutants have longer disc perimeters (E) owing to their crescent or horseshoe shapes. The mean disc radius of curvature (a measure of doming) is also markedly reduced in MBPKD mutants as compared wild-type dG-GFP (delta-giardin-GFP) tagged strains (F). Triple asterisks indicate p values < 0.001 in t-tests.

Morpholino knockdowns or stable CRISPRi knockdowns of DAP16343 display similar mutant phenotypes with the almost complete loss of disc doming (Figure 3A) with phenotypic penetrance ranging from 0-40% (morpholino knockdown) to up to 90% (CRISPRi) dependent on guide RNA position [17]. Here the aberrant disc phenotype of the MBPquadKO is stable and 100% penetrant (Figure 3B,C), with the majority of disc mutants possessing a crescent-moon phenotype that lacks an overlap zone and roughly 30% of disc material. The remainder of cells have either a horseshoe shaped or a half moon shaped disc phenotype, representing slightly more or slightly less disc present (Figure 3B). On average the total area of mutant discs (40 µm^2^ ) is only slightly less than wild type (44 µm) (Figure 3D) yet due to the “open” non-overlapping disc, the perimeters of mutant discs are roughly 10 µm greater that wild type. To quantify disc doming in mutants, we quantified the angle of repose in MBPKD mutants. Despite the lower penetrance of the phenotype in transient morpholino knockdowns (∼40%) this angle was dramatically lower on average as compared wild type. This mutant angle of repose is similar to that of chilled, detached trophozoites (Figure 2B).

The complete phenotypic penetrance of the MBPquadKO mutant also facilitated imaging of the mutant discs at high resolution (TEM, Figure 4). We observed disrupted and irregular spacing of the disc MT spiral, with noticeable gaps in the crossbridges linking adjacent microribbons (Figure 4D,E,F). In cross-section, it is clear that while microribbons remain trilaminar [12], they have irregular heights and the spacing between MTs is disrupted (Figure 4F). Some microribbons appear to be bifurcated or fused (Figure 4F). The lack of the overlap zone is clear in the MBPquadKO and the ends of the disc spiral MT abruptly end (arrow, Figure 4D). Microribbons in the ventral groove region also appear disordered and unevenly spaced relative to wild-type (arrow, Figure 4E).

**Figure 4:**
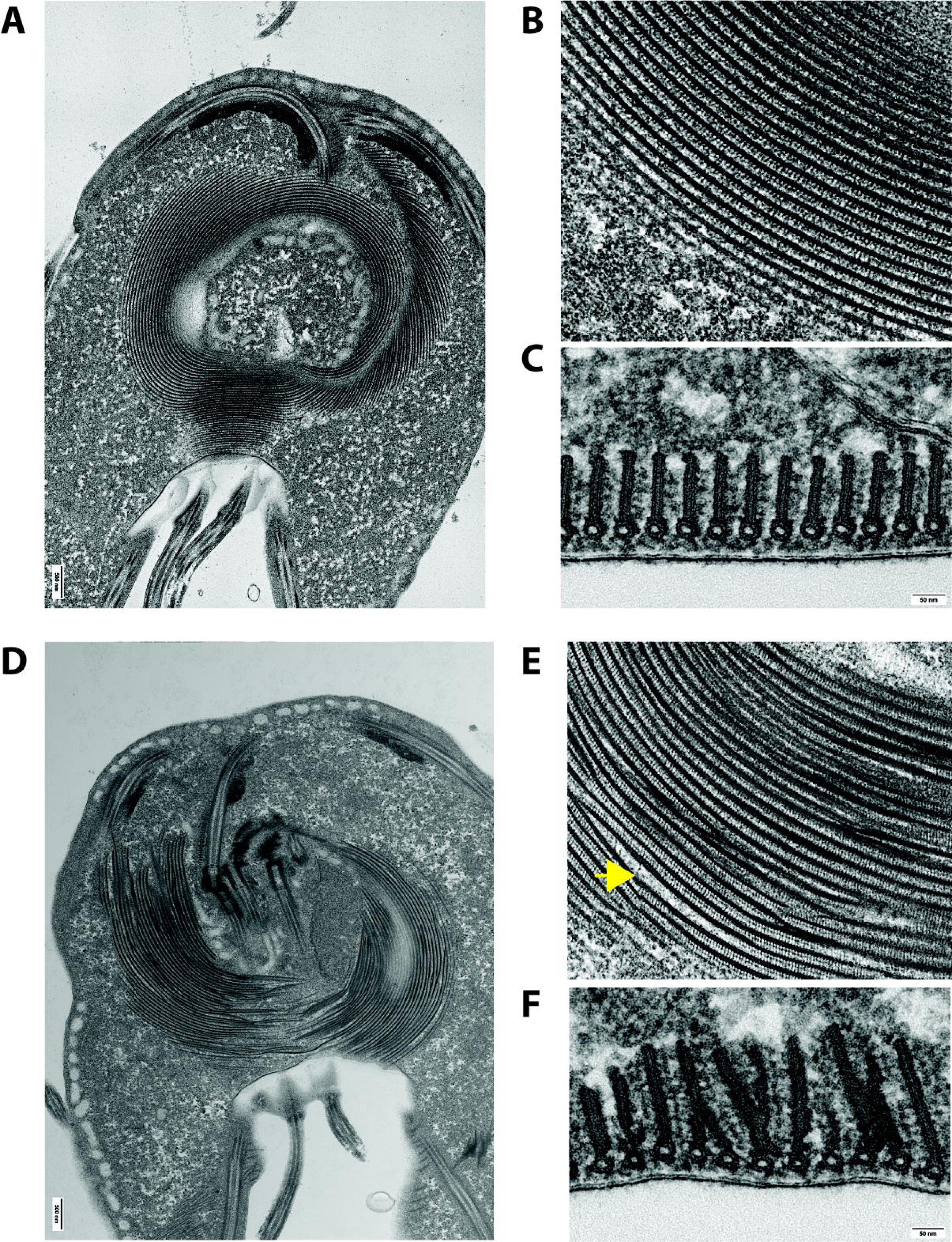
MBPquad KO mutants have severe structural and spacing defects in microribbon-crossbridge complexes. Transmission electron microscopy with sectioning of wild-type WBC6 whole trophozoite cells highlight the spiral array of microtubules (MTs) (A) that scaffold additional complexes such as the microribbon-crossbridge complexes thought to maintain regular spacing of microribbons (MRs) and repetitive crossbridges (B). In C, the cross section demonstrates the repetitive and regular sized MR-CB complex emphasizing the microtubules (MT) oriented into the page, vertical microribbons (MR) and crossbridges (CB). In D, MBPquadKO mutant disc emphasize the irregular and incomplete microtubules and MR-CB complexes. Panel E shows the irregular spacing of the MR-CB complex often with large regions lacking crossbridges (arrow). In vertical sections of MBPquadKO mutants, the MR have markedly irregular heights and are often fused and irregularly spaced (F).

### Disc KO doming mutants have decreased attachment under flow and lack the ability to form a proper lateral crest and bare area contacts with the surface

We have previously defined stages I to IV of *Giardia* attachment based on surface contacts using TIRFM [20], and have shown that transient, morpholino- and stable CRISPRi-based knockdowns of MBP have a reduced ability to attach to surfaces as quantified by shear force flow assays and centrifuge-based normal force attachment assays [20]. We have also previously shown that morpholino knockdown results in trophozoites that lack proper surface contacts and are unable to properly attach to surfaces [20]. Here, we examined surface contacts of attached, membrane stained, wild-type and MBPquadKO cells (Figure 5A). Most wild-type cells (93%) exhibited the hallmarks of Stage IV attachment—a complete lateral crest “seal” and normal surface contacts of the bare area (the region in the center of the disc with many vesicles, but devoid of MTs). The remainder were at Stage III (normal lateral crest seal but lacking bare area contact). In contrast, only 3% of the completely penetrant MBPquadKO cells created a normal lateral crest “seal”(Figure 5A). Due to the abnormal disc structure of MBPquadKO trophozoites, lateral crest and bare area contacts for the majority of attached MBPquadKO mutants were unlike anything previously observed for wild-type, necessitating two additional categories, Stage III-LC (no lateral crest seal and no bare area contact, 22%) and Stage IV+B (a partial lateral crest seal with exaggerated bare area contact; 75%).

**Figure 5:**
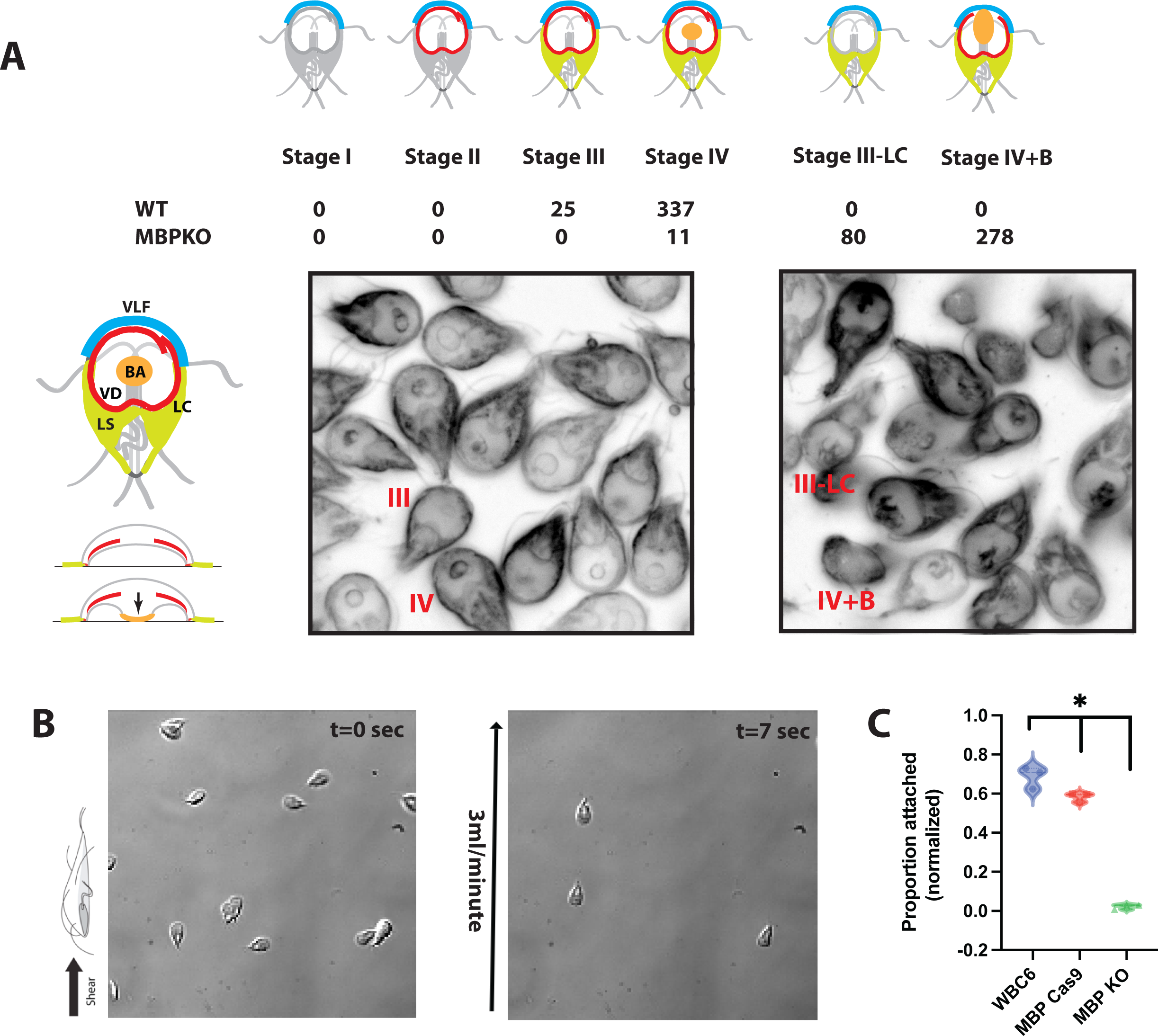
MBPquadKO mutants have aberrant surface contacts and an inability to resist shear stress under flow. In panel A, surface contacts associated with various early stages of trophozoite attachment (stages I-IV, [20]) are scored in both MBPquadKO and wild-type strains. MBPquadKO mutants lack canonical surface contacts, and instead have two additional attachment stage categories: stage III-LC which is similar to stage III but lacks lateral crest contacts, and stage IV+B which is defined by aberrant bare area contacts connecting to the lateral crest. Shear stress assays using 3ml/minute flow (B) highlight the severe defects in the ability of MBPquadKO mutants to resist shear stress, specifically, the reduced proportion of attached cells following 20 seconds of shear stress as compared to the MBPCas9 plasmid alone strain and the WT strain (C). N = 3 separately cultured flasks, tested individually. Triple asterisks indicate p values < 0.01 in t-tests.

To evaluate shear forces of attachment, we used a novel quantitative shear stress assay that employs time lapse imaging of attached parasites in commercial (Ibidi) flow chambers (Methods). MBPquadKO, MBP Cas9 control plasmid (MBP gRNA only), and wild-type strains were attached in chambers before challenging with a flow rate of 3 ml/min that causes shear forces detaching most wild-type trophozoites (Figure 5B). Close to 80% of the attached wild type parasites resisted 3ml/min of flow (∼4dyn/cm2), and a majority of the parasites containing the MBP Cas9 control plasmid resisted the same flow (60%). In contrast, the MBPquadKO mutants had very little resistance to 3ml/min of flow (Figure 5B), with less than 5 % of parasites remaining attached.

### MBPquadKO mutants lack the ability to significantly impact epithelial barrier breakdown in human gastrointestinal organoid-derived monolayers (ODM)

2D human intestinal organoid-derived monolayers (ODMs) mature after about 9 days of culture, when they reach a height of approximately 20 μm, and the cells exhibit the typical polarized shape of enterocytes, with microvilli and well-developed cellular junctions [36]. At this stage, ODMs exhibit a progressive increase in trans-epithelial electrical resistance (TEER), which stabilizes at around 250 Ω · cm2 after 9 days of culture. TEER is a proxy for barrier function integrity which decreases significantly in the *Giardia* infections of ODMs [36].

To study effects of wild-type and mutant *Giardia* trophozoites on degradation of barrier function, ODMs were infected with both wild-type trophozoites and two MBPquadKO clones (MBPKO1 and MBPKO2) and mock infections (no *Giardia*) at multiplicities of infection (MOI) ranging from 1 to 10, with saturation of trophozoites at the ODM surface at MOI 10 (Figure 6A). In wild-type, normalized TEER decreases in a dose- and time-dependent manner after 24 hours of infection and peaks at 48-54 hours (Figure 6A and 6B). In contrast, both MBPquadKO clones had significantly more normalized TEER as compared to wild-type control infections.

**Figure 6:**
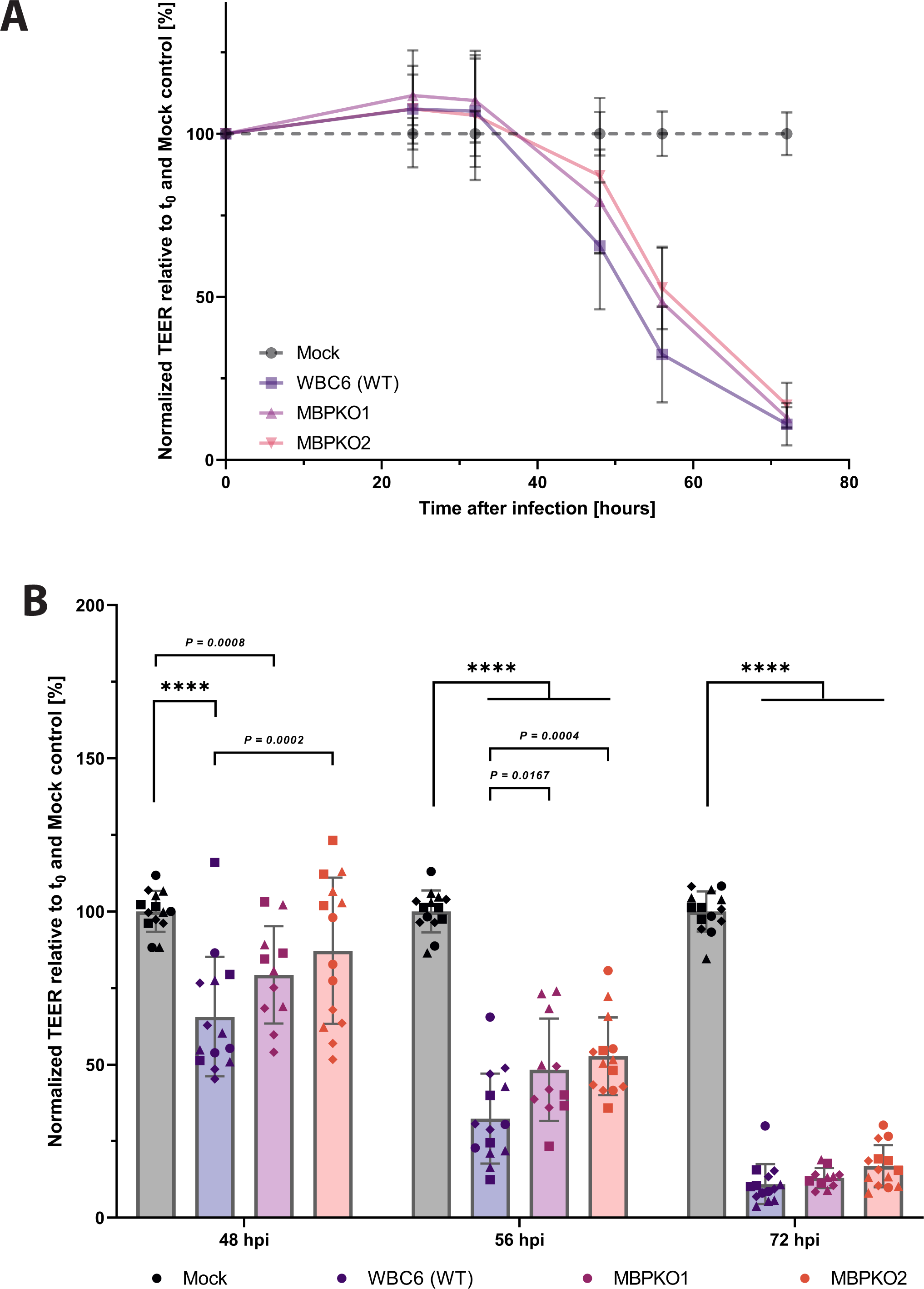
MBPKO attachment mutants have a significantly reduced impact on enterocyte barrier degradation. TEER measurements of ODM-based infections of wild-type (WBC6) trophozoites and two MBPquadKO disc mutant clones (MBPKO1 and MBPKO2) are compared with mock (no *Giardia*) infections in an 80 hour time period (A). In B, relative TEER measurements are compared at representative time points (0, 48, 56, 72 hours post infection (hpi), with significant differences indicated by T-tests (****).

## Discussion

The elaborate ventral disc architecture consists of curved, regularly spaced singlet MTs that scaffold additional protein complexes such as the microribbon-crossbridge, or MR-CB, complex. The identities of DAPs comprising the MR-CB complex are largely unknown, yet at least three SF-assemblins (beta-giardin, delta-giardin, and SALP1) are microribbon proteins [11]. The microribbons extend roughly 400 nm into the cytoplasm and are connected by repetitive crossbridge structures that presumably maintain roughly 50 nm spacing between adjacent MTs of the spiral. While the MR-CB complex has been associated with both structural [13] [14] and functional roles [15] in the ventral disc, direct evidence for either role of this complex is generally lacking.

### New genetic approaches transform the study of Giardia biology and disc-mediated host-parasite interactions

By combining disc proteomics with protein-tagging and imaging approaches [16,18], our lab has identified close to 90 disc-associated proteins. However, predicting the functions of the majority of these DAPs is problematic, as most lack any homology to known MT-binding proteins, with roughly 30% of DAPs lacking any homology to known proteins outside *Giardia*, and another 30% of DAPs containing only ankyrin-repeat domains. Even with new machine learning approaches such as AlphaFold, which is based on known protein structures from model organisms [37], only 12 of 87 DAPS have the majority of their structures predicted with overall high model confidence (>70) using the per-residue confidence metric (pLDDT) outside of conserved protein-protein interaction motifs (Supplemental Table 1). Thus, both the role of specific DAPs in generating the overall curved and domed cup-like structure, and the potential contributions of specific DAPs to disc-mediated attachment remain to be determined.

Our lab and others have recently used new molecular genetic strategies to create transient or stable knockdowns and have identified DAPs essential for disc hyperstability [18], disc biogenesis, and/or attachment [20]. One such DAP – DAP16343 (MBP, median body protein) – was initially described using immunostaining to be a component of the median body [38]; it was later found by our lab to instead be an abundant and essential component of the disc with occasional median body localization [33]. DAP16343 is a 100 kDa DAP that lacks homology to proteins in species other than *Giardia* and consists primarily of three coiled coil domains [38]. Coiled coil proteins are known to influence organelle architecture and can propagate conformational changes or function as molecular spacers [39]. Within the disc, DAP16343 localizes to the disc body, disc margin, and the overlap zone, the region of overlap between the upper and lower portions of the disc (Figure 1A-C and [11,16]) that appear to contact one another.

Initial morpholino-based translational knockdown [20] and later CRISPRi-based transcriptional knockdown of DAP16343, or “MBP” [17] provided the first molecular genetic evidence that this single DAP is essential for both ventral disc ultrastructure and proper attachment under hydrodynamic flow. While they are powerful genetic approaches, these knockdown strategies are limited by the degree of phenotypic penetrance and, in the case of morpholinos, by their transience—morpholinos are diluted each generation and phenotypes are lost within 1-2 days in *Giardia* [17]. CRISPRi-based knockdown mutants are stable *in vitro*, but contain episomal plasmids that must be maintained by positive selection [17]. This makes them unsuitable for evaluating host-parasite interactions because the long periods of cultivation of parasites off selection that are required when performing *in vivo* infection studies in animal models (7 to 14 days) [40] or new human intestinal organoid models (48 hours or more) [36] promote plasmid loss and subsequent loss of the mutant phenotype.

Here we used a CRISPR-based knockout strategy based on our CRISPRi system [17] with new positive selectable markers to create a DAP16343 quadruple null mutant in the double diploid *Giardia* (Methods). The DAP16343 KO mutant has dramatic attachment defects and resolves prior issues of phenotypic transience, incomplete penetrance, or reliance on antibiotic selection to maintain mutant phenotypes. Our new knockout approach permits structural and functional evaluations of mutant attachment phenotypes on both biological and non-biological surfaces, and for the first time, we were able to use a new organoid model to provide direct evidence for a role of disc-mediated attachment in causing the cellular defects associated with *Giardia* infections.

### MBP is required for proper MR-CB complex formation and the overlapped, domed disc architecture

The ventral disc MBP knockout mutants have a characteristic horseshoe or half-moon disc phenotype and lack both disc doming and the disc overlap zone region [20]. As compared to our prior knockdowns of DAP16343 with morpholinos or CRISPRi, the MBPquadKO phenotype is 100% penetrant. While roughly 10% of discs resemble flattened horseshoes or half-moons, over 85% have more severe defects and resemble crescents (Figure 3). Despite the open discs and lack of the overlap zone, the total area of the disc in the MBPquadKO mutant is comparable to wild type. Mutant trophozoites also have reduced surface contacts and cannot resist shear forces. Other aspects of the MBPquadKO mutant resembles wild-type *Giardia,* including flagellar beating amplitude and frequency and ATP generation (data not shown).

As seen in cross-section with TEM (Figure 4), the microribbon-crossbridge complex is severely malformed in MBPquadKO mutants. MRs lack regular spacing and have non-uniform heights. Disc MT spacing is also disrupted and irregular. In total, the disrupted MR-CBs, the lack of an overlap zone, and the lack of disc doming support the prediction that MBP is an essential disc component likely associated with the MR-CB complex itself. Because depletion of MBP results in mis-assembled MR-CB complexes at the local level of MTs, the overall flattened and crescent-shaped disc argues that proper MR-CB complexes are required to create the domed disc architecture required for attachment. The lack of an overlap zone also confirms an essential contribution of MBP in linking the upper and lower regions of the disc. The lack of the overlap zone also results in an aberrant or incomplete disc margin, which is the perimeter of the spiral MT array and contacts the ventral surface, yet includes a gap at the point of the overlap zone [9].

With respect to disc architecture, DAPs could directly (or indirectly) contribute to nucleation of the spiral disc MTs, stabilization of MT plus and minus ends, or creation of the overall curvature and doming of the disc (Brown et al., 2016; Schwartz et al., 2012). Our knowledge of the assembly and biogenesis of the ventral disc, and especially the assembly of the MR-CB complexes, is limited, nor do we understand the mechanisms coordinating ventral disc microtubule and microribbon polymerization and the targeting of DAPs to specific regions [11]. This phenotype also implies that MBP contributes to MR-CB complex biogenesis, but is not necessary for partial disc assembly. Our analysis of mutant phenotypes also supports assembly of the MR-CB complexes relatively late in disc biogenesis after MR-CB complexes associate with the disc MT spiral. Because the overall MT curvature remains despite the crescent shape of aberrant MBPquadKO discs, we predict that additional DAPs govern MT curvature of the spiral [13,14].

### A domed disc is a necessary component of suction-based attachment to inert surfaces

*Giardia* can attach to surfaces regardless of surface treatment (PEG, Teflon) [41], which supports a suction-based attachment model for attachment to inert surfaces. Attachment is also readily reversible, and trophozoites prefer to attach to flat surfaces rather than to pillars [8]. This indifference to surface chemistry may help explain *Giardia*’s zoonotic potential by allowing the parasite to attach to a variety of mammalian intestinal environments despite varying luminal chemistry. Yet hydrodynamic suction requires a means to not only generate forces for attachment but also a means to maintain a negative pressure underneath the disc. The oft-cited hydrodynamic model [42] posited that the continued beating of the ventral flagella is required both to generate and maintain hydrodynamic suction and the associated the negative pressure differential underneath the disc. Through the genetic disruption of motility of all flagella and the ventral flagella specifically [38], we previously showed that flagellar motility is not sufficient for trophozoites to maintain attachment to surfaces once attachment has been initiated. Flagellar motility could be necessary, however, for the initiation of attachment, including intestinal site recognition and orientation of the disc toward the attachment surface. A negative pressure differential for suction-based attachment to inert surfaces could be generated by a variety of means, including dynamic movements of the structure of the disc itself.

Regardless of the mechanism of force generation for attachment, we confirm here that the domed disc architecture is required for parasites to attach to inert surfaces and withstand hydrodynamic current, as the flattened and opened discs of the MBPquadKO mutant have a drastically reduced ability to resist fluid flow. As we have reported previously, cell surface contacts beyond those of the ventral disc (e.g., lateral shield, ventrolateral flange, etc) are also observed during attachment [8], and on inert or biological surfaces, these contacts likely promote cellular adhesion in addition to disc-mediated attachment (Figure 5).

Most experimental evaluation of attachment still does not distinguish among mechanistic differences associated with *attaching* as compared to already *attached* trophozoites [40]. Attachment of wild type parasites involves a stepwise progression through four distinct attachment stages, which we previously defined using TIRFM imaging of wild type trophozoites attaching to glass [9]. Trophozoites first encounter the attachment plane and then skim along the surface, making membrane/surface contacts with the anterior portion of the ventrolateral flange. As parasites attach, the membrane around the disc perimeter forms a seal that begins with anterior contacts at the ventrolateral flange proceeds around the disc perimeter and ends at the ventral groove region. The presence of this lateral crest seal, along with contacts of the bare area with the surface, demarks the transition from attaching trophozoites to attached trophozoites that are resistant to flow. We predict that MBPquadKO mutants are unable to progress through these four early stages of attachment due to an inability form a suction-based seal defined by the characteristic bare area contacts [17].

Attached trophozoites are commonly detached from the inert surfaces of laboratory growth vessels such as tubes or plates by incubation for 15-30 minutes at 4°C [21]. As demonstrate here, cold discs are flat; all other treatments including paraformaldehyde fixation, MT rugs, or detergent extraction resulted in discs that remained domed as they are in wild-type trophozoites (Figure 2). MBPquadKO mutants, however, have severe attachment defects, and the flat discs phenocopy discs incubated on ice that detach. The disruptions of the cell surface contacts and lateral crest seal caused by open, flattened discs in the MBP knockout are reminiscent of phenotypes observed with morpholino or CRISPRi based MBP knockdowns, but are more severe and more penetrant. Taken together, these multiple lines of evidence support the domed conformation of the disc is required for attachment to surfaces, regardless of the attachment force. We predict that the rigidity and doming of the disc allows for space underneath the disc to maintain a compartment with a negative pressure differential relative to the outside medium on inert surfaces.

### Disc doming and parasite attachment are required for epithelial barrier breakdown in organoids

*Giardia* infections have been associated with various detrimental effects to the host epithelium, including villus shortening, decreased trans-epithelial electrical resistance (TEER), increased permeability, altered ion and glucose transport, and increased apoptosis. Any direct or indirect impact of *Giardia’s* extracellular attachment to observed cellular pathology seen in in *Giardia* infections has been largely overlooked in investigations of pathogenic mechanisms of giardiasis [9]. Our current understanding of disease mechanisms has instead relied heavily on the use of immortalized cancer cell lines, such as Caco-2 cells derived from colon carcinoma, to investigate the effects of *Giardia* infection on host tissues. Based on this work, it is proposed that *Giardia*’s secretion of proteases leads to the activation of caspase-3 in the host, resulting in the disruption of tight junctions (TJ) and caspase-dependent apoptosis. Prior work has yielded conflicting results, however, likely due to differences in tight junction composition between cell lines and that of the small intestine, which is the primary site of *Giardia* infection. Additionally, Caco-2 cell lines do not respond well to the growth medium required for *Giardia* infections [36].

The recent development of primary organoids derived from small intestinal stem cells [6,36]has opened new avenues for understanding *Giardia* pathogenesis, particularly considering that inflammation does not necessarily play a significant role in the disease. In brief, 3D organoid cultures are derived from human duodenal biopsy specimens and used to establish a 2-dimensional transwell system known as organoid-derived monolayers (ODMs). *Giardia* infection of ODMs leads to a gradual disruption of the epithelial barrier, which is influenced by the duration of infection and the parasite load. Infection also triggers changes in ODM gene expression related to ion transport and the structure of tight junctions. Recent work in ODM models also point to alternative mechansims by which *Giardia* induces cellular pathologies. In particular, MLCK and caspase inhibitors used in ODM models to limit cysteine protease or caspase activity do not prevent the breakdown of trans-epithelial electrical resistance (TEER) caused by the parasite or the increased occurrence of apoptosis.

Here for the first time, we use 100% penetrant disc KO mutants with significant attachment and structural defects (Figures 3-5) in ODM-based infections to determine if attachment is necessary for barrier breakdown in TEER assays (Figure 6). We provide direct evidence linking parasite attachment to cellular pathologies in the host epithelium (Figure 6). Both MBPquadKO clones have significantly limited ability to induce barrier breakdowns and underscore the necessary role parasite attachment in inducing complete barrier breakdown in this model. Other factors are likely involved as we do not see a complete lack of barrier breakdown, yet the close and intimate association provided by extracellular attachment does impact host cellular pathology and likely contributes to observed systemic symptoms of giardiasis.

## Conclusion

Attachment by the complex ventral disc is a hallmark feature of *Giardia*’s adaptation to life in close association with the host gastrointestinal epithelium. While we demonstrate here that the domed disc architecture is necessary for attachment to the host, there are clearly undiscovered or uninvestigated disc components and mechanisms of attachment required for trophozoites to grip and attach to host enterocyte microvilli. The discovery here of a role for attachment in contributing to barrier breakdown also opens up new potential mechanisms for *Giardia*-host interactions. This discovery highlights the multifaceted nature of *Giardia*-host interactions beyond indirect metabolic interactions. Lastly, the new availability of powerful genetic approaches and *in vitro* ODM models holds promise for subsequent *in vivo* studies with stable *Giardia* attachment mutants to illuminate the intimate interactions of *Giardia’s* ventral disc during attachment to the host.

## Acknowledgements

This work was supported by and a R01AI077571 award to SCD. The human MCF10A cell line was a kind gift from Nont Kosaisawe and Dr. John Albeck (Dept. MCB, UC Davis). We graciously thank the UC Davis MCB Microscopy Core for helpful advice on the SDC and SIM microscopes. We also thank Shane McInally (Brandeis) for providing helpful feedback on manuscript drafts.

## Supplemental Figures and Tables

**Supplemental Figure 1: Antibiotic cassette design for HDR templates**

**Supplemental Figure 2: Test of antibiotic sensitivities and antibiotic cassettes in Wt Giardia**

**Supplemental Figure 3: Design strategy of antibiotic cassettes and homology arms used for quadruple KO**

**Supplemental Figure 4: PCR-based screening of quad MBPKO and clone lines**

**Supplemental Figure 5: 3D projection of image stack of disc curvature of delta-giardin GFP strain to intestinal cell line**

**Supp Table 1: Alpha fold summaries of DAPs (GiardiaDB)**

## Notes

### Competing Interest Statement

The authors have declared no competing interest.

### Summary of Updates

revised supplemental data and fixed reference problems

