## Supplementary figures and images for "The domed architecture of *Giardia*’s ventral disc is necessary for attachment and host pathogenesis"

### Supplemental Figure 3

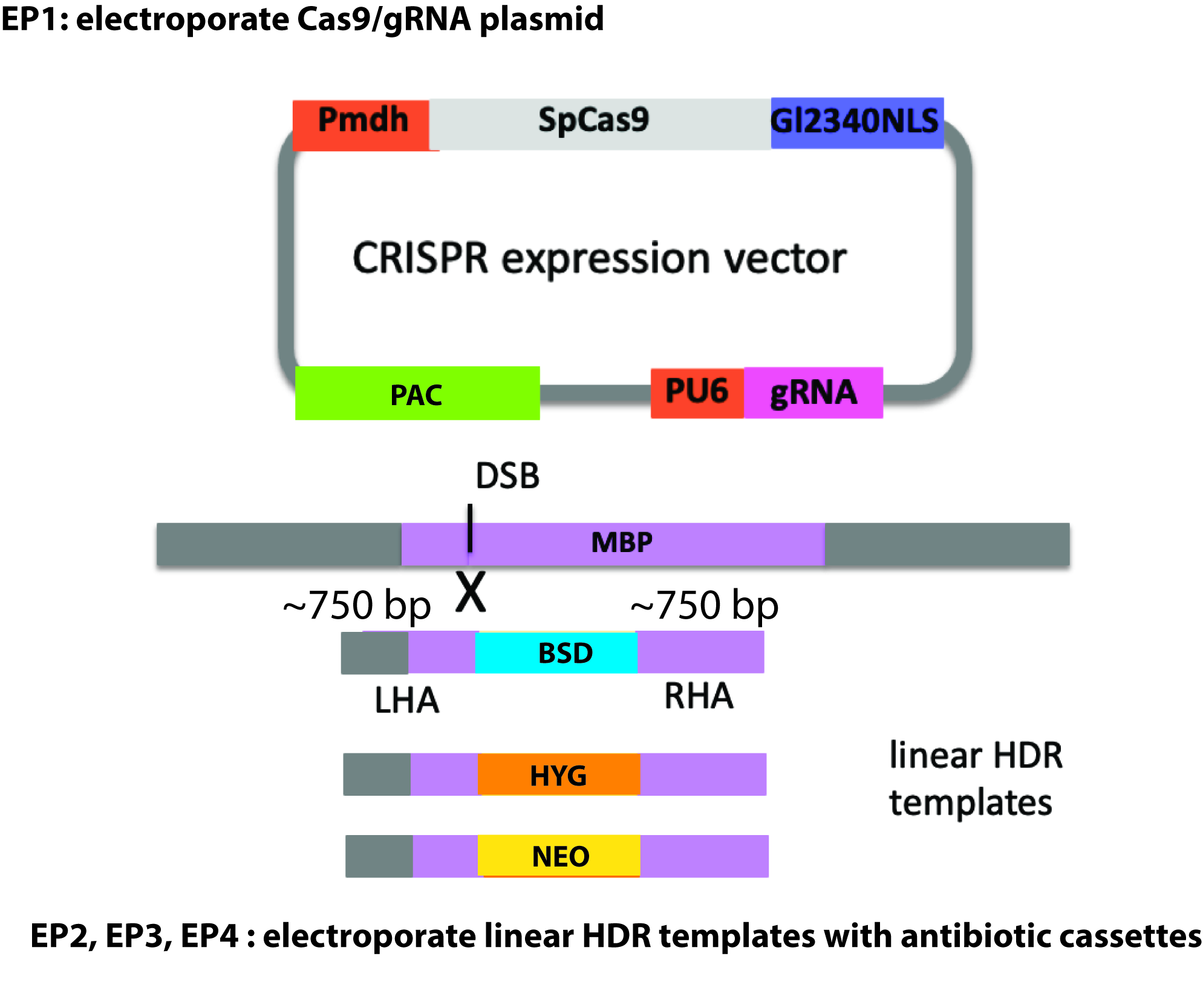
