## Supplemental Figure 4 for "The domed architecture of *Giardia*’s ventral disc is necessary for attachment and host pathogenesis"

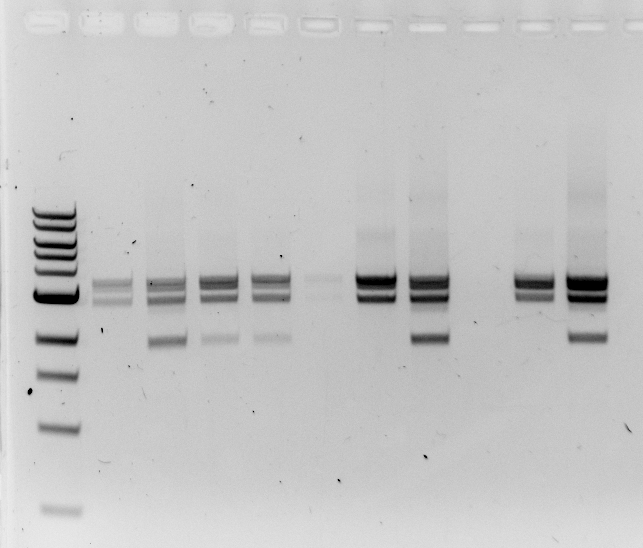

Dilution to extinction of the well #9 population yielded MBP both triple and quadruple KO clones:

wt

+Bsd

+Neo

+Hyg

3.0 kb

2.0 kb

1.5 kb

4.0 kb

9A

9B

9C

9D

9E

9F

9G

9H

9I

9J

*

*

*

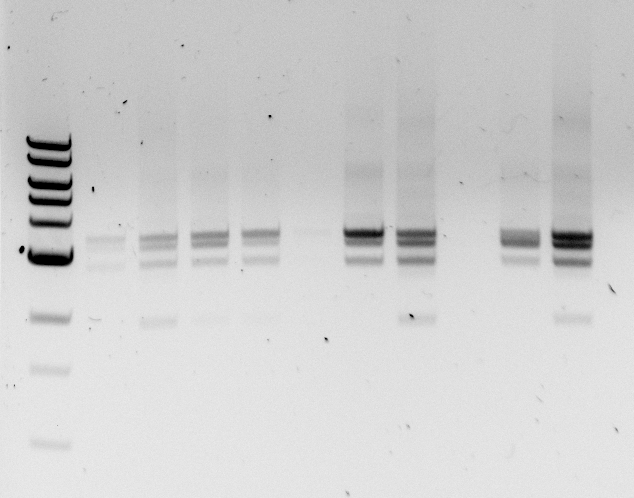

4.0 kb

3.0 kb

2.0 kb

wt

+Bsd

+Neo

+Hyg

Further electrophoresis

of the same gel resolved

the Hyg and Neo bands.

Clones 9A, 9F and 9I

lack wild type copies

of MBP
